# The bacterial hitchhiker’s guide to COI: Universal primer-based COI capture probes fail to exclude bacterial DNA, but 16S capture leaves metazoa behind

**DOI:** 10.1101/2021.11.28.470224

**Authors:** Sanni Hintikka, Jeanette E.L. Carlsson, Jens Carlsson

## Abstract

Environmental DNA (eDNA) metabarcoding from water samples has, in recent years, shown great promise for biodiversity monitoring. However, universal primers targeting the *cytochrome oxidase I (COI)* marker gene popular in metazoan studies have displayed high levels of nontarget amplification. To date, enrichment methods bypassing amplification have not been able to match the detection levels of conventional metabarcoding. This study evaluated the use of universal metabarcoding primers as capture probes to either isolate target DNA or to remove nontarget DNA, prior to amplification, by using biotinylated versions of universal metazoan and bacterial barcoding primers, namely metazoan *COI* (mlCOIintF) and bacterial *16S* (515F). Additionally, each step of the protocol was assessed by amplifying for both metazoan *COI* (mlCOIintF/jgHCO2198) and bacterial *16S* (515F/806R) to investigate the effect on the metazoan and bacterial communities. Bacterial read abundance increased significantly in response to the captures (*COI* library), while the quality of the captured DNA was also improved. The metazoan-based probe captured bacterial DNA in a range that was also amplifiable with the *16S* primers, demonstrating the ability of universal capture probes to isolate larger fragments of DNA from eDNA. Although the use of the tested *COI* probe cannot be recommended for metazoan enrichment, based on the experimental results, the concept of capturing longer fragments could be applied to metazoan metabarcoding. By using a truly conserved site without a high-level taxonomic resolution as a target for capture, it may be possible to isolate DNA fragments large enough to span over a nearby barcoding region (e.g., *COI*), which can then be processed through a conventional metabarcoding-by-amplification protocol.

## INTRODUCTION

The use of environmental DNA (eDNA) metabarcoding for monitoring aquatic species has steadily increased during the past decade (Thomsen et al. 2012, Port et al. 2016, O’Donnell et al. 2017, Sigsgaard et al. 2017, Cilleros et al. 2019, Jeunen et al. 2020). In many cases, the aim of a metabarcoding effort is to estimate the number of different species or taxa present in a body of water, without having to physically observe the organisms. Although many studies have shown that eDNA metabarcoding is more sensitive than traditional monitoring methods (Valentini et al. 2016, Sard et al. 2019), others have pointed out that choices made in the laboratory and during the bioinformatics analyses can have large impacts on the inferred species lists and downstream ecological analyses (Ficetola et al. 2014, Evans et al. 2017).

Analysing eDNA present in highly diverse marine samples is associated with well-known challenges, especially when targeting metazoans. For example, the proportion of available eukaryotic DNA in a marine environmental sample can be overrun by orders of magnitude more of genetic material from microbial origins (Stat et al. 2017, Zafeiropoulos et al. 2021). Additionally, a recent study comparing the effectiveness of metabarcoding water eDNA samples against bulk samples has demonstrated large discrepancies in the biodiversity obtained between these two methods (Hajibabaei et al. 2019), noting especially the inability of eDNA metabarcoding to detect key indicator species. That said, neither is necessarily an issue when species-specific primers are used on eDNA samples, as positive detections can be made from very low concentrations (Biggs et al. 2015, Furlan et al. 2016, Gargan et al. 2021). Recently, less universal primers were developed targeting macroinvertebrates while specifically excluding nontarget groups (Leese et al. 2021). While these primers capture mostly target DNA and thus can give a deeper picture of the community present, they also show more primer bias than compatible universal primers, sometimes struggling to detect specific species or even groups. Many metabarcoding studies target the mitochondrial gene *cytochrome oxidase I (COI),* the classical eumetazoan barcoding gene, which due to its high rate of evolution give it good interspecific resolution. Yet, this complicates the development of universal primers aimed at amplifying *COI* for a broad range of taxa in a single metabarcoding effort (Collins et al. 2019). These universal primers for *COI* are often highly degenerated, which can lead to high levels of nontarget amplification, and therefore also amplify nontarget DNA (especially when little target DNA is present in the sample), as well as introduce primer bias during amplification (Elbrecht and Leese 2015, Collins et al. 2019, Zafeiropoulos et al. 2021). For marine eDNA samples it is not uncommon to have over 50% of the resultant Actual Sequence Variants (ASVs) or molecular operational taxonomic units (mOTUs) assigned to bacterial origins due to *COI*-like bacterial genes co-amplifying with the degenerate *COI* primers (Zafeiropoulos et al. 2021).

To address the issues of primer bias, enrichment methods that bypass the amplification step altogether have been investigated in the context of metabarcoding (Zhou et al. 2013, Dowle et al. 2016, Mariac et al. 2018, Wilcox et al. 2018). On bulk samples, whether using differential centrifugation to isolate mitochondrial DNA (Zhou et al. 2013) or employing 20,000 capture probes designed based on a database of target taxa (Dowle et al. 2016), the enrichment results in detection rates that are comparable to or better than conventional PCR-based methods, with the added benefit of good correlation of returned sequence abundance to the relative biomass or abundance of species in the sample. To avoid the cost and technical challenges of synthesising multiple probes via PCR in-house (Maggia et al. 2017), Mariac et al. (2018) used a single capture probe synthesised in-house that was designed to be equidistant to the fish species found in the Amazon basin. Captures were done on ichthyoplankton samples with results showing good correlation to real species frequencies as discovered by morphological identifications. Despite the encouraging results, few have tested capture approaches on aquatic eDNA samples where low levels of target DNA is the norm rather than the exception. The first proof of concept for enrichment from eDNA samples came from Wilcox et al. (2018), who synthesised capture probes via PCR from 40 target taxa. Although they found a good correlation of true abundance to sequences per taxa, the sensitivity of the method was deemed much lower than that of contemporary PCR-based metabarcoding.

This study wanted to take advantage of the universality of existing primers used in metabarcoding and apply them to the capture process. The theory behind this approach relates to the varying probabilities of primers amplifying a specific target. Based on Kebschull and Zador (2015), if the probability of a template to get amplified by a primer varies only during the first few cycles of PCR (and not after), the difference between a “lucky” and “unlucky” template can be up to 4-fold by the end of 25 cycles. Applying their theory into metabarcoding, then, having a much higher proportion of amplifiable non-target DNA can potentially counteract the higher probability of target templates being amplified, throughout the PCR cycles, purely due to numbers of each type of available template. However, if the target templates that have a higher probability of being amplified with the primer, are isolated after the first cycle, the relative proportions of target and amplifiable non-target DNA should be evened out to an extent.

To test this theory, biotinylated versions of two common metabarcoding primers for metazoan and bacterial metabarcoding were used in an attempt to isolate either nontarget or target templates prior to the typical amplification step. Sequencing for both target and nontarget markers allowed for an assessment of the effect of the capture protocol steps on both communities.

## MATERIALS AND METHODS

The eDNA samples chosen for this study were collected in July 2019 as part of a larger sampling effort characterising habitat connectivity (results reported elsewhere). The samples used here were from two subsites of a coral reef habitat (one on a small reef outcrop and one on the reef proper) known as Banco Capiro in Tela, Honduras (15°51’48.6”N 87°29’42.9”W), following the sampling and filtering protocols as outlined in Supplementary File 1 – Section 1. A total of ten eDNA samples (five from each subsite) were extracted using Qiagen Blood and Tissue kit as per manufacturers protocol and eluted in 100 μl of AE buffer. To investigate the effect of the capture protocol on the diversity detected from these eDNA samples, the experiment was designed to track changes at each step of the protocol in both metazoan and bacterial communities by building sequencing libraries targeting the metazoan *cytochrome oxidase subunit I (COI)* and the bacterial *16S* small subunit rRNA *(16S).*

## THE EXPERIMENTAL DESIGN

To assess changes in the detected community, a total of six steps of the capture protocols were amplified and sequenced for each of the eDNA samples. The experimental design is visualised in Figure 1. Firstly, the untreated eDNA was amplified to provide a baseline against which the rest of the steps could be compared (step ID: raw_eDNA). To evaluate the effect of a basic bead clean, step 2 involved a clean with AMPure XP beads at a 1:1 ratio, using 20 μl of the eDNA sample (step ID: AMPure). Steps 3 and 4 involve the capture of DNA with a *COI* probe, to assess the community that can be detected from the *COI* captured product (step ID: *COI*_capt.) and from what is left behind after the capture (step ID: *COI*_eluent). The same was done for steps 5 and 6, where the eDNA sample was subjected to a capture with a 16S probe, and both the captured product (step ID: 16S_capt.) and the eluent (step ID: 16S_eluent) were amplified to assess the effect on the communities detected. Two libraries were constructed for sequencing; one for *COI* and one for bacterial 16S, to better quantify and assess the effect of the captures in the target and nontarget groups.

**Figure 1:**
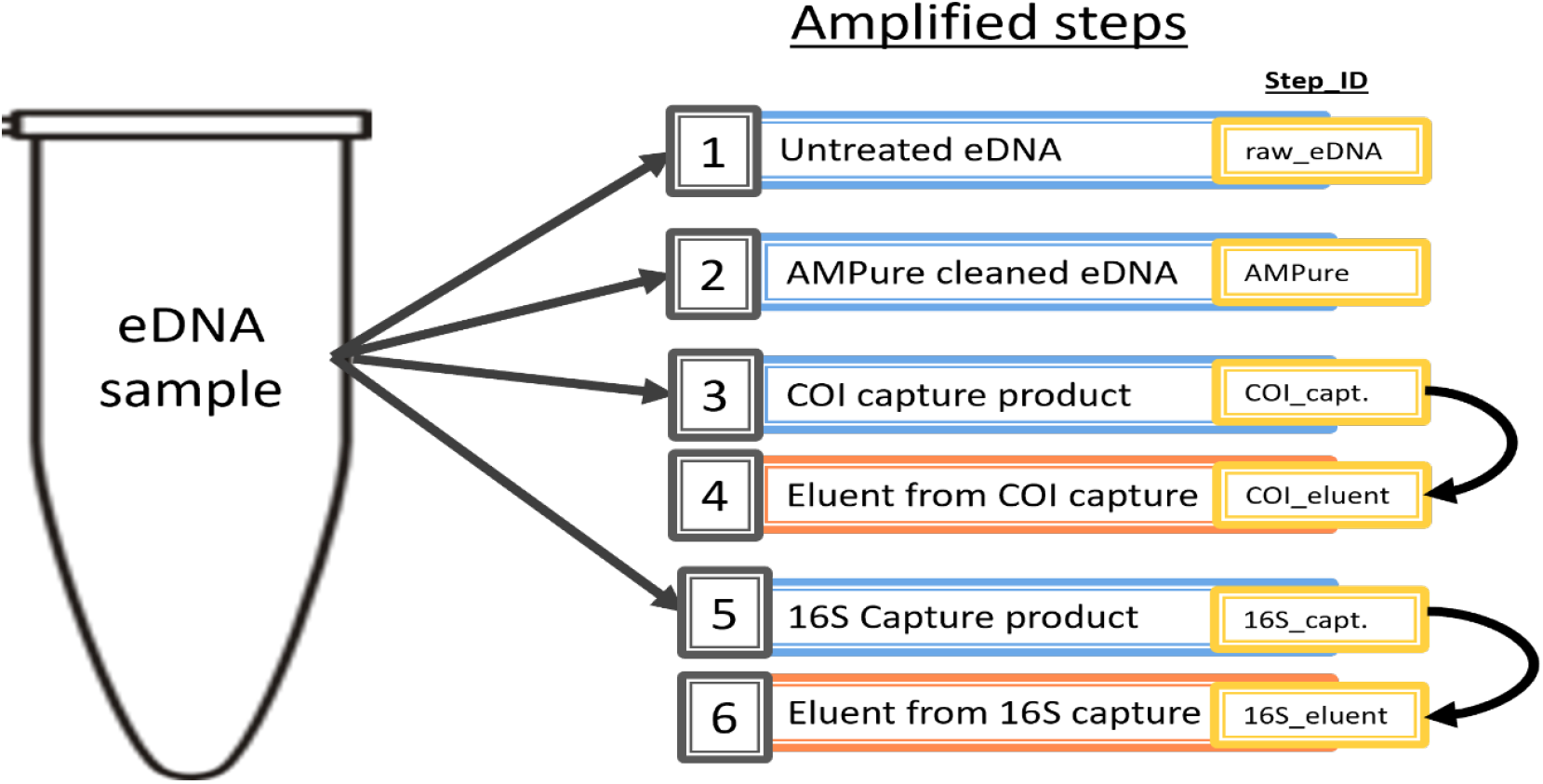
Flowchart of targeted capture experimental design: Experimental design to assess changes in community detections through the targeted capture protocol (eDNA sample n = 10). The numbered steps detail the type of template used in PCR to amplify both metazoan *COI* and bacterial *16S.* Untreated eDNA (i.e., raw_eDNA) refers to template taken directly from eDNA extract. AMPure clean refers to template from eDNA extract that was cleaned with AMPure magnetic beads. Steps 3 and 6 (capture steps) refer to the product of the targeted capture with either *COI* and *16S* probe, resp., and are explained in more detail in main text. Steps 4 and 6 are templates obtained from the eluents of steps 3 and 5, respectively, rather than from direct processing of the eDNA sample. The IDs used for each step in the figures throughout this study are shown in the yellow boxes.

### Targeted capture method – Capture probes

For the captures targeting the *COI* region, a biotinylated version of forward primer mICOIintF (referred to as *COI* probe; Leray et al. 2013) was used (5’-biotin-GGW ACW GGW TGA ACW GTW TAY CCY CC-3’). For the captures targeting bacterial *16S*, a biotinylated version of forward primer 515F (referred to as 16S probe; Earth Microbiome Project (EMP)) was used (5’-biotin-GTG YCA GCM GCC GCG GTA A – 3’).

### Targeted capture method – Hybridisation and bead capture

For both *COI* and *16S* capture reactions, hybridisation was performed using 20 μl of eDNA sample and 20 μl of hybridisation buffer (12 μl of SSC (20X), 0.2 μl of SDS (10%), 1.2 μl of BSA (10 mg/ml), 1.6 μl of probe (*COI* or 16S, 10 μM), 5 μl of ddH_2_O). Both hybridisation reactions were first incubated for 10 minutes in 95°C followed by 24 h in 58°C or 63°C for the *COI* and 16S probes, respectively. Then, the reactions were allowed to cool to room temperature before processing with streptavidin coated magnetic beads (DynaBeads© M270).

Streptavidin binds biotin and allows for the target gene/region to be pulled out of the samples with the annealed biotinylated probes using an external magnet. The beads were prepared as per manufacturer’s instructions, by washing stock beads (concentration 10 μg/μl) thrice with 2X binding and washing buffer (2X B&W buffer: 10 mM Tris-HCl (pH 7.5), 1 mM EDTA, 2 M NaCl), and resuspending in twice the stock volume of 1X B&W buffer, to a final concentration of 5 μg/μl. For the capture, a 1:1 ratio of beads to sample was deemed most appropriate (pers. comm. with DynaBeads© manufacturer) to achieve the optimal 3D spatial configuration of beads in the reaction and to maximise their biotin binding ability. Therefore, 35 μl of the hybridised product and 35 μl of the prepared beads were mixed in new 0.5 μl PCR tubes by pipetting 10 times and incubated for 30 minutes at room temperature with gentle rotation. After incubation, the tubes were placed on a magnetic plate for 3 min to collect the beads with their bound probe-target complex on the side of the tubes. The eluent (approximately 70 μl) from each capture was transferred to new 1.5 ml Eppendorf tubes, and the magnetically captured products were washed three times with 1X B&W buffer before being resuspended in 20 μl ddH_2_O (to match the original eDNA volume used for the capture). A visualisation of the targeted bead capture process is provided in Figure 2.

**Figure 2:**
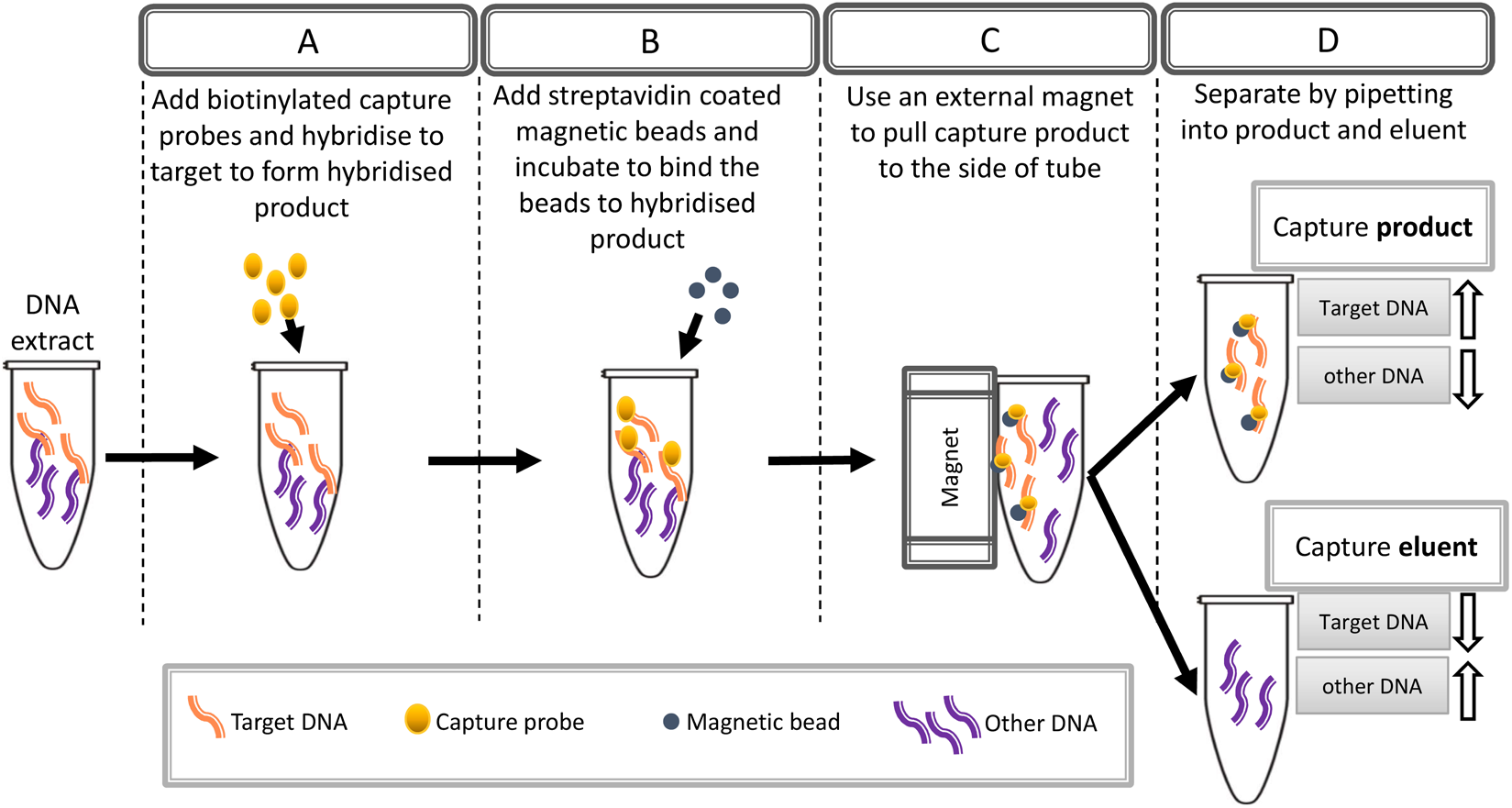
Flowchart describing the stages of a targeted bead capture protocol.

The eluents were further processed with ammonium acetate precipitation to reduce, after pilot testing showed evidence of potential PCR inhibition, likely due to the high salt concentration of the buffers used (pilot data not shown). First, for each 70 μl eluent, 5.6 μl of ammonium acetate (7.5 M) and 160 μl of cold 100% EtOH was added, and the tubes centrifuged at 12 400 rcf or 30 min. Then, the liquid was discarded (if no visible pellet, approx. 3-5 μl was left in tubes) and another 100 μl of cold 70% EtOH was added, and the tubes centrifuged at 12 400 rcf for another 15 minutes. Again, the liquid was discarded, but if no pellet was visible approx. 3-5 μl was left in the tube and let evaporate on a thermal block set to 50°C. Finally, the precipitated DNA was re-eluted in 20 μl of ddH_2_O, to match the volume of the original eDNA sample used for the capture reaction.

To account for any contamination that might have occurred during sample processing, protocol controls (PC) were included throughout the process. Essentially, for every step where DNA was added or manipulated, an additional sample with ddH_2_O in place of DNA was included and brought through the rest of the protocol (i.e., for a capture of *COI*, one sample with water was also put through the capture process, with the capture product of the water sample acting as the capture PC, and the eluent from that used as the eluent PC).

## AMPLIFICATION AND LIBRARY PREPARATION

To multiplex the samples, a combination of ten and seven uniquely tagged forward and reverse primers (respectively, for details see Supplementary_file_1 – Section 2: Table S1) (*COI*: 5’ – 6bp tag – mlCOIintF – 3’ / 5’ – 6bp tag – jgHCO2198 – 3’ (Leray et al. 2013); *16S*: 5’ – 6bp tag – 515F – 3’ / 5’ – 6bp tag – 806R – 3’ (EMP)) were used to amplify an approx. 313 bp section of the mitochondrial *cytochrome oxidase I (COI)* gene and the approx. 250 bp hypervariable V4 region of the bacterial *16S* gene, respectively. A different tag combination was used for each step of the protocol and each eDNA sample used (for details see Suppl. file – Metadata.csv, columns F_tag and R_tag), and PCR reactions were run in triplicate. Each 30 μl reaction consisted of 2 μl template DNA, 1.2 μl of each forward and reverse primers (10 μM), 1.2 μl dNTP’s, 1.2 μl BSA (10 mg/ml), 0.12 μl KAPA Taq Polymerase, 3.0 μl KAPA Taq Buffer A and 20.08 μl ddH_2_O. The thermocycling profile for *COI* amplification consisted of an initial denaturation step of 5 min at 95°C, followed by 40 cycles of 10s at 95°C, 30s at 46°C and 60s at 72°C, and a final elongation step of 10 minutes at 72°C. For *16S* amplification, the thermal conditions were an initial denaturation step of 3 min at 94°C, followed by 35 cycles of 45s at 94°C, 60s at 50°C and 90s at 72°C, and a final elongation step of 10 minutes at 72°C. The PCR replicates were then pooled for each tag combination, excess primers and nucleotides were removed using ExoSAP-IT™ (ThermoFisher Scientific) according to manufacturer’s instructions, the replicate pools were quantified and finally pooled in equimolar amounts into the *COI* and *16S* libraries. The final library preparation (PCR-free sequencing adapter ligation) and sequencing using NovaSeq 250PE technology was undertaken by Novogene Europe.

## BIOINFORMATIC PROCESSING

Bioinformatic processing was done for the *16S* and *COI* libraries separately. The quality of the raw reads was first assessed with FastQC (v0.11.9, Andrews 2010). Due to the library preparation method leading to a mixed orientation of reads in the sequencing output (both forward and reverse primers are possible in both paired end raw files, i.e., in R1 and R2), for each library, demultiplexing was performed twice using CUTADAPT (Martin 2011), once for each orientation. Additionally, to account for the tag sequences being the same for both forward and reverse primers, the full tag plus primer sequences were used for demultiplexing (instead of only the 6bp tag sequences) with a maximum two errors allowed. Full demultiplexing commands are provided in Supplementary_file_1 – Section 3.

The rest of the bioinformatic processing was done using the DADA2 pipeline (v1.12.1, Callahan et al. 2016), with each orientation processed separately to avoid mixing of error models as per developer’s recommendation. The following describes the steps taken for each orientation. First, minimum read length was set to 100bp, and both read directions (R1 and R2) were set to truncate at 200bp (function: *filterAndTrim;* parameters: truncLen=c(200,200), minLen=100, maxN=0, maxEE=c(2,2), truncQ=2). Then, to denoise the demultiplexed reads into Actual Sequence Variants (ASVs) DADA2 estimates the sequencing error rates from the demultiplexed data. The pipeline was modified to accommodate the use of Illumina NovaSeq 6000 data instead of MiSeq (which the pipeline was developed for) by enforcing monotonised decreasing error rates after the error estimations, as recommended by the pipeline developers. Finally, the paired-end reads were merged across all samples, filtered to 253 bp and 313 bp length for the 16S and *COI* libraries, respectively, and chimeras were removed (details of read abundance at each step of the pipeline are available in suppl. file Metadata_all.csv).

The resulting ASVs were then taxonomically assigned using the RDP Naïve Bayesian classifier method (Wang et al. 2007). This classifier uses kmers to find the best hit in a reference database and assigns a bootstrap confidence value at each taxonomic level. For the *16S* data, the DADA2 function *assignTaxonomy* was used with the SILVA version 132 trained database, which covers Bacteria, Archaea and Eukaryotic taxa for the *16S* small subunit rRNA locus. For the *COI* ASVs, the RDP assignment tool was used outside of DADA2, with a trained database of *COI* references covering Bacteria, Archaea and Eukaryotic taxa (Porter and Hajibabaei 2018). For each *16S* and *COI* assignment, a bootstrap threshold of 0.5 was applied across all taxonomic levels, as recommended by Wang et al. (2007). A further curation step to remove spurious ASVs and cluster similar ASVs together was done with the LULU algorithm (Frøslev et al. 2017).

Control correction to remove potential contaminant ASVs was first applied using the negative controls (no-template controls and extraction blanks) and a relative abundance threshold of 10%; if an ASV found in a control sample had a relative abundance of >10% out of the total abundance for that ASV across all samples, it was discarded (as in Antich et al. (2021)). The same approach was applied with the protocol controls (PCs) but applying the correction only to samples of the same protocol step (i.e., PC2 = *COI* capture: ASV relative abundances calculated across *COI* capture samples only).

## STATISTICAL TESTS

All statistical testing and data analyses were performed in R (v4.0.3) using packages *phyloseq* (v1.34.0, McMurdie and Holmes (2013)) and *vegan* (v2.5.7, Oksanen et al. (2009)). Changes in the reads obtained through the bioinformatic pipeline were assessed as total reads and as a percentage of reads that passed from demultiplexing through control correction. Kruskal-Wallis nonparametric tests to assess variation of means were done across all template types in each library (*COI* and *16S*), and pairwise Wilcoxon rank sum tests were carried out comparing the raw eDNA template to all other template types. After the passing reads statistics were computed, all samples were rarefied without replacement to sample depth of 30 000 reads (minimum sample depth was 30 111 reads).

For the *COI* library, relative read abundance per kingdom was plotted for each template type, and both total reads and ASV richness were plotted as boxplots for reads assigned to groups Metazoa, Bacteria and Unassigned. Kruskal-Wallis tests and paired Wilcoxon rank sum tests with raw eDNA as reference group were calculated for each of the investigated groups (Metazoa, Bacteria and Unassigned). Furthermore, the Unassigned ASVs were taxonomically placed using the Dark mAtteR iNvestigator tool (DARN, Zafeiropoulos et al. (2021)) in order to gain a better understanding of the ASVs that could not be assigned using the RDP taxonomic assignment method. The placed Unassigned ASVs were qualitatively assessed using kronaplot visualisations from DARN.

For the *16S* library, relative read abundances of the top phyla making up approx. 99% of the total reads were plotted for each template type, and both total reads and ASV richness were plotted as boxplots for reads assigned to Proteobacteria, Cyanobacteria and Bacteroidetes, as they were found to be the top three phyla. Kruskal-Wallis tests and paired Wilcoxon rank sum tests with raw eDNA as reference group were calculated for each of the top three phyla.

To assess whether the capture protocol influenced template quality, total reads and richness of ASVs that had at minimum a family level assignment were examined using Kruskal-Wallis and paired Wilcoxon tests for both libraries.

## RESULTS

The individual steps for which results are presented for are outlined in Figure 1. To summarise briefly, “raw eDNA” refers to templates taken directly from eDNA extracts, “*COI* capture” and “16S capture” refer to templates that were captured from the raw eDNA extract using either *COI* or 16S capture probes (resp.), and “*COI* eluent” and “16S eluent” refer to what is left behind, after the capture product has been removed with either *COI* or 16S capture probes (resp.). All steps were amplified with *COI* and 16S primers to build two sequence libraries, and each step included ten samples.

For the two libraries combined, a total 19 419 054 raw reads were returned from the NovaSeq sequencing effort (raw sequence files available at European Nucleotide Archive (ENA) under project accession PRJEB49001), of which 9 728 354 passed through demultiplexing, denoising and control correction. For the *COI* library, the raw eDNA samples yielded the highest number of total reads passing the bioinformatics pipeline, but in the 16S library, only the *COI* eluent showed significantly less reads than raw eDNA passing bioinformatics and control correction (Fig. 3A). No significant differences in the proportion of reads per sample that passed all bioinformatics steps were observed between the raw eDNA versus any other step of the protocol, indicating no loss in overall sequence quality was observed (Fig. 3B). Overall richness of ASVs in the *COI* library was not affected by the capture protocols steps, but the 16S library showed significant decreases of ASV richness after the AMPure clean, as well as both eluents and the 16S capture templates (Fig. 3C)

**Figure 3:**
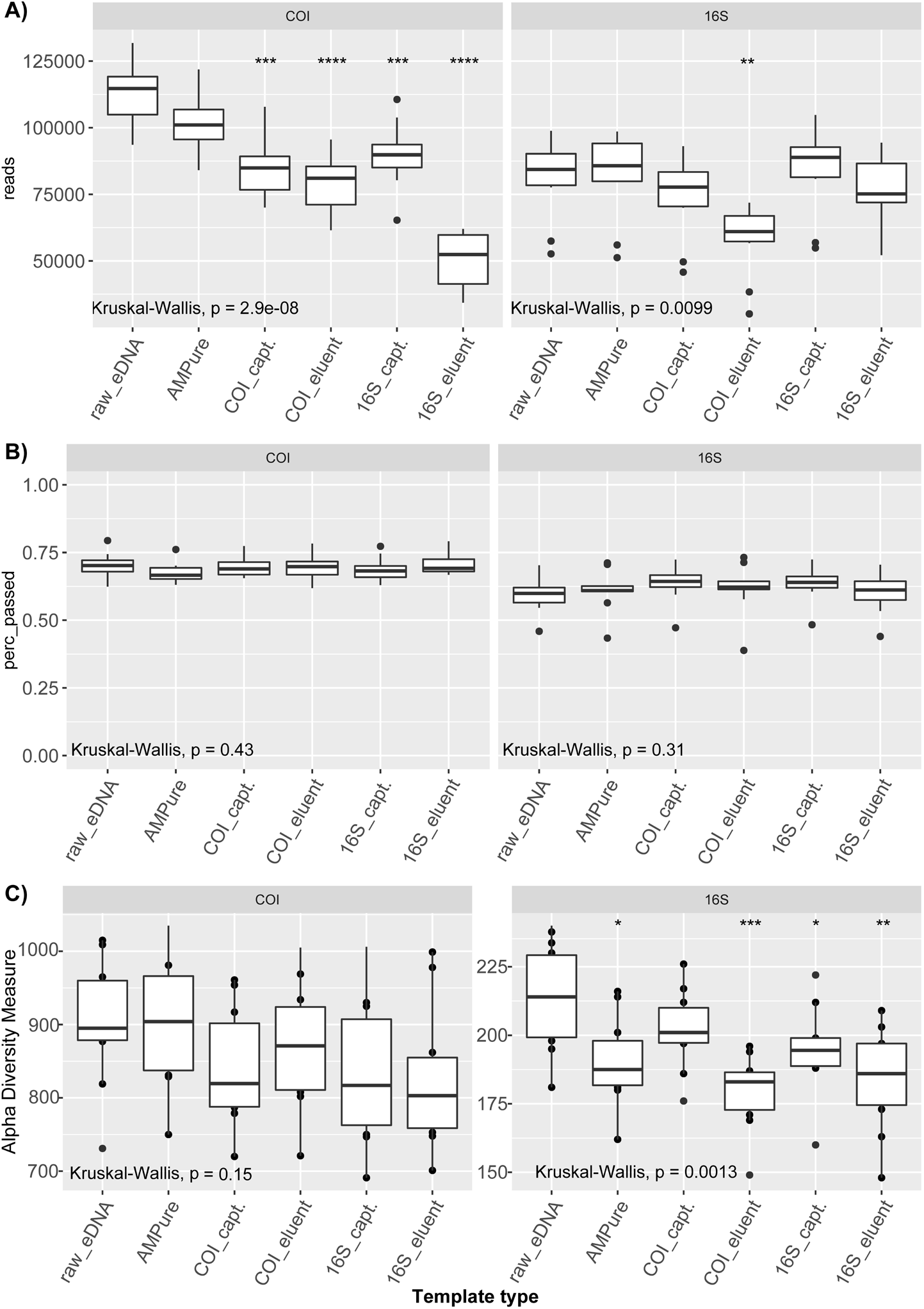
Reads and ASVs through bioinformatics and processing. Total reads (A) that passed the bioinformatics pipeline (denoising, length filtering, merging, chimera removal and control correction) and proportion of demultiplexed reads (B) that passed the bioinformatics pipeline (denoising, length filtering, merging, chimera removal and control correction). C) The total richness of ASVs in each library after rarefying to even sequencing depth of 30 000 reads/sample. Box limits depict the 25%-75% interquartile ranges with the horizontal line showing the median value (n=10 for each template type). The whiskers extend to the upper and lower quartiles, and outliers are shown as points. Kruskal-Wallis non-parametric test p-values are shown for each group, and significant Wilcoxon pairwise test results between the raw eDNA samples and each of the other template types are depicted with asterisks (* = p<0.05, ** = p < 0.01, *** = p < 0.001, **** = p < 0.0001). For details of template type, please refer to Fig. 5.1. NOTE: The y-axes in A do not start at 0 to accommodate the large scales, and in C the y-axes of *COI* and 16S are on different scales for a better visualisation.

## *COI* LIBRARY

In total, 2 074 ASVs and 1.8M reads were included in the analyses of the *COI* library after rarefying the samples to a read depth of 30 000. It was hypothesised that both relative read count and ASV richness for metazoans would be increased with the templates from *COI* captures or eluents of the *16S* captures, as in each case the expectation is that target template availability is increased in relation to the nontarget. However, Unassigned ASVs dominated the results originating from each template type (Fig. 4). Additionally, a slight increase was observed in the relative abundance of both bacterial and metazoan reads when using either the *COI* or 16S capture products as template.

**Figure 4:**
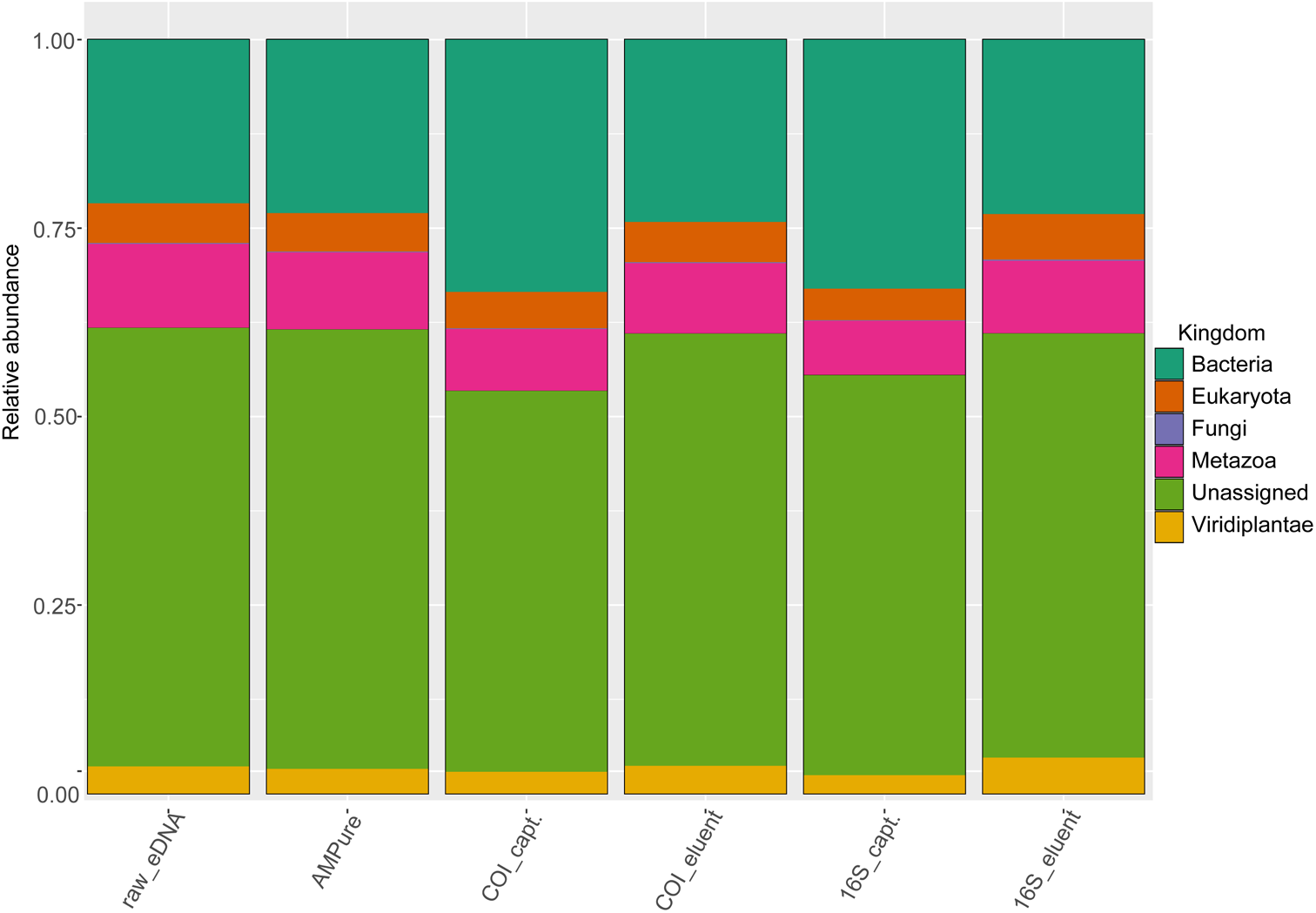
*COI* library relative abundance. Relative abundance of kingdoms for each originating template type.

Total metazoan reads were observably reduced in both capture products when compared to the raw eDNA samples, but this was only statistically significant for the 16S captures (Fig. 5A). The ASV richness of metazoans was significantly reduced for both capture products (*COI*_capt. and *16S*_capt.) (Fig. 5A). On the other hand, bacterial reads showed a significantly higher abundance for both the *COI* and 16S captures when compared to the raw eDNA but did not display any statistically significant changes in ASV richness, despite overall evidence of significant variation (Fig. 5B). The unassigned reads exhibited a significant drop in read abundance for the *COI* capture and 16S capture template types, but no significant differences were observed in the richness of unassigned ASVs (Fig. 5C).

**Figure 5:**
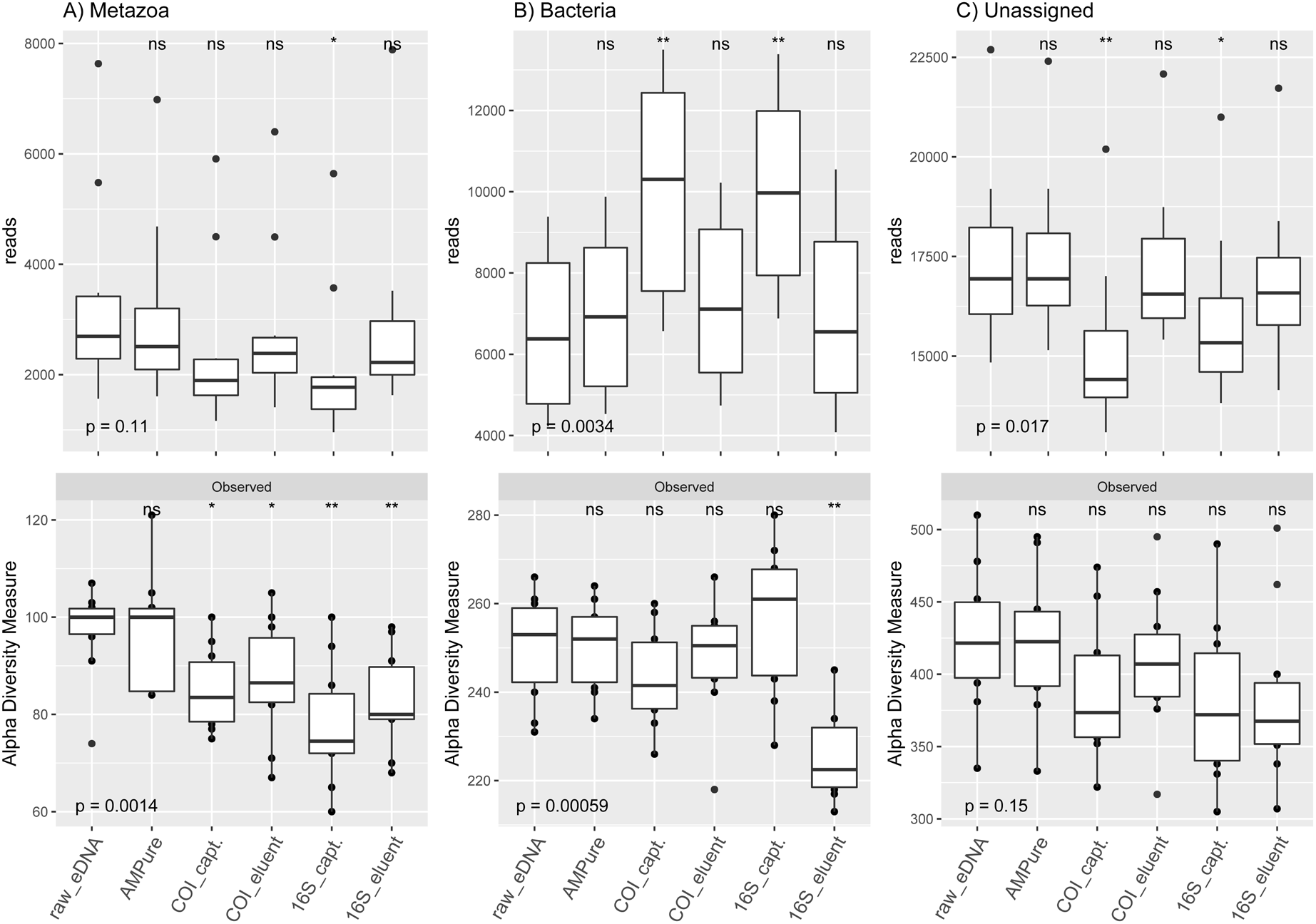
*COI* read abundance and ASV richness. Total reads (top) and ASV richness (bottom) for Metazoa (A), Bacteria (B) and Unassigned ASVs of the *COI* library. On the x-axis are the different template types used from each step of the targeted capture protocol. The p-values at the bottom show the result of the Kruskal-Wallis test for the whole group, while significant results of Wilcoxon pairwise test between reference group raw_eDNA and all other template types are depicted with asterisks. (Significance levels: * = p < 0.05, ** = p < 0.01). The boxplot limits stretch over the interquartile range, with the horizontal line signifying the median value, and the whiskers reaching to the upper and lower limits. Outliers are shown as points. NOTE: y-axes do not begin at 0 to accommodate the scale differences for the three phyla and to provide a clearer visualisation.

The Unassigned ASVs that were further taxonomically placed with DARN did not seem to vary to any considerable extent between the different template types (see suppl. file Captures_uA_krona.html) in terms of broad taxonomic groupings. The majority were placed within Eukaryota (between 76% (*16S* capture) and 78% (raw eDNA)), of which between 42% to 46% were further placed within the Haptophyta, a protist group of phototrophic and mainly planktonic marine organisms that are currently known from 330 described species. Of the eukaryotically placed ASVs, another approx. 30% for each template type did not receive any deeper placement than Eukaryota, while approx 20-25% were divided between various small and/or microbial groups such as Chlorophyta (green algae), Basidiomycota (fungi), Dictyochophyceae (heterokont algae) and Cercozoa (mostly heterotrophic protozoa).

## *16S* LIBRARY

The 16S library contained 406 ASVs in 1.8M reads after rarefying to even sequencing depth (30 000 reads) without replacement. Unsurprisingly, bacterial reads dominated the 16S library, with between 98.2% and 99.1 % of reads assigned to Bacteria for each template type (Supplementary File 1: Fig. S1). A total of six phyla made up 98.6% of all the reads in the rarefied dataset. The most abundant phylum across all template types was Proteobacteria, followed by Cyanobacteria and Bacteroidetes (Fig. 6, Supplementary File 1: Fig. S2). Among the top six phyla was also one archaeal group, Euryarchaeota. No differences in relative abundances of phyla between the template types were observed in the 16S library, except in the 16S eluent where the relative abundance of Cyanobacteria was visibly increased, while Proteobacteria were reduced.

**Figure 6:**
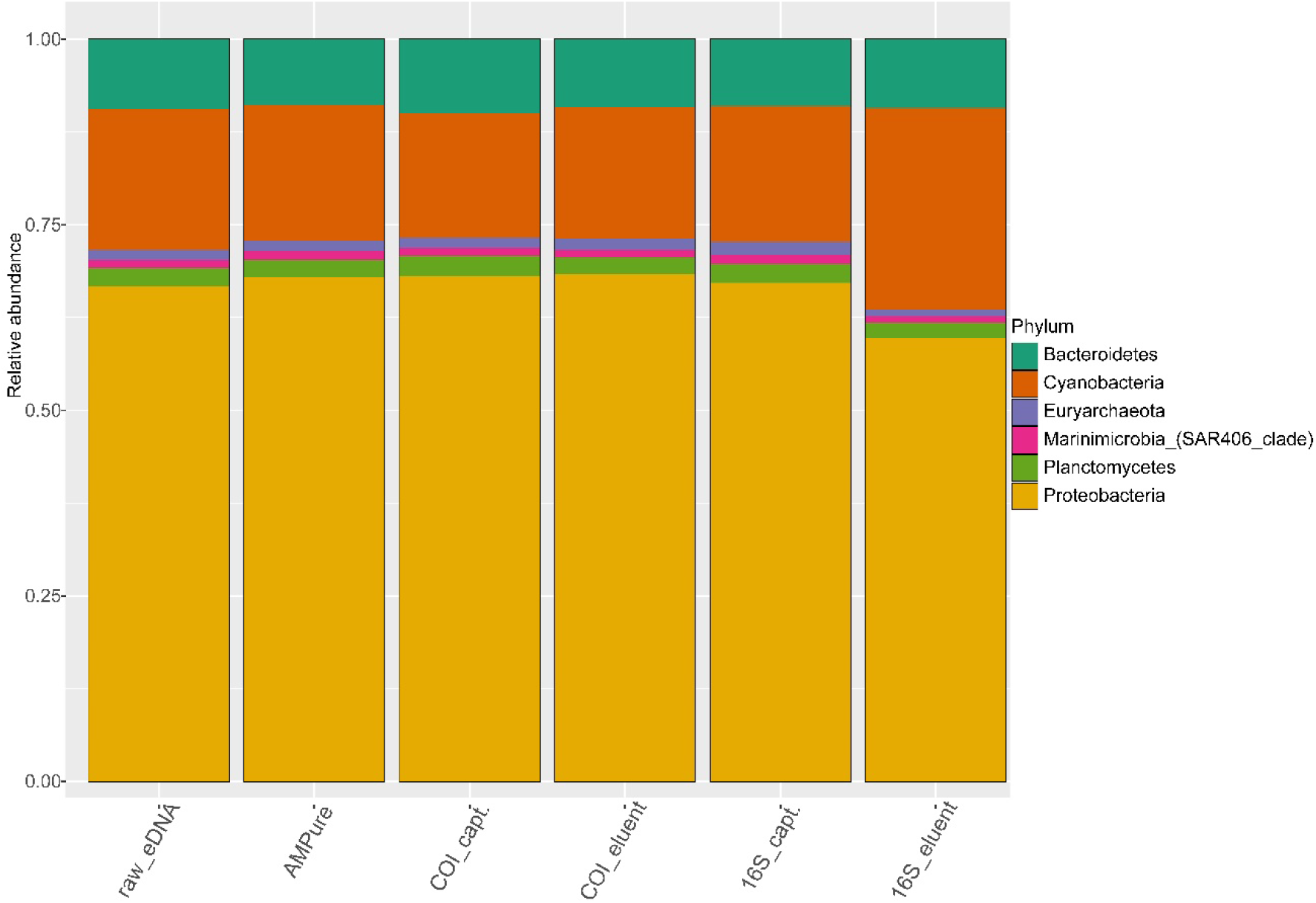
*16S* library relative abundance. Relative abundance of six most abundant phyla detected in the *16S* library with each template type.

Neither Proteobacteria nor Bacteroidetes showed statistically significant changes in the reads obtained throughout the capture protocol, but the Cyanobacteria phylum displayed a significant increase in total reads in the 16S eluent when compared to raw eDNA (Fig. 7 A-C). ASV richness varied significantly in all the top three phyla. For the dominant phylum Proteobacteria, all template types showed a decrease in richness when compared to the raw eDNA (Fig. 7A). Cyanobacteria exhibited a drop in richness for the AMPure cleaned templates, as well as the 16S eluent, despite the increase in reads (Fig. 7B). Meanwhile, richness was reduced for the Bacteroidetes phylum for template types of *COI* eluent, 16S capture and 16S eluent. It should be noted that the overall richness for the most dominant phylum was approx. four to five times higher than the two others used in detailed analysis.

**Figure 7:**
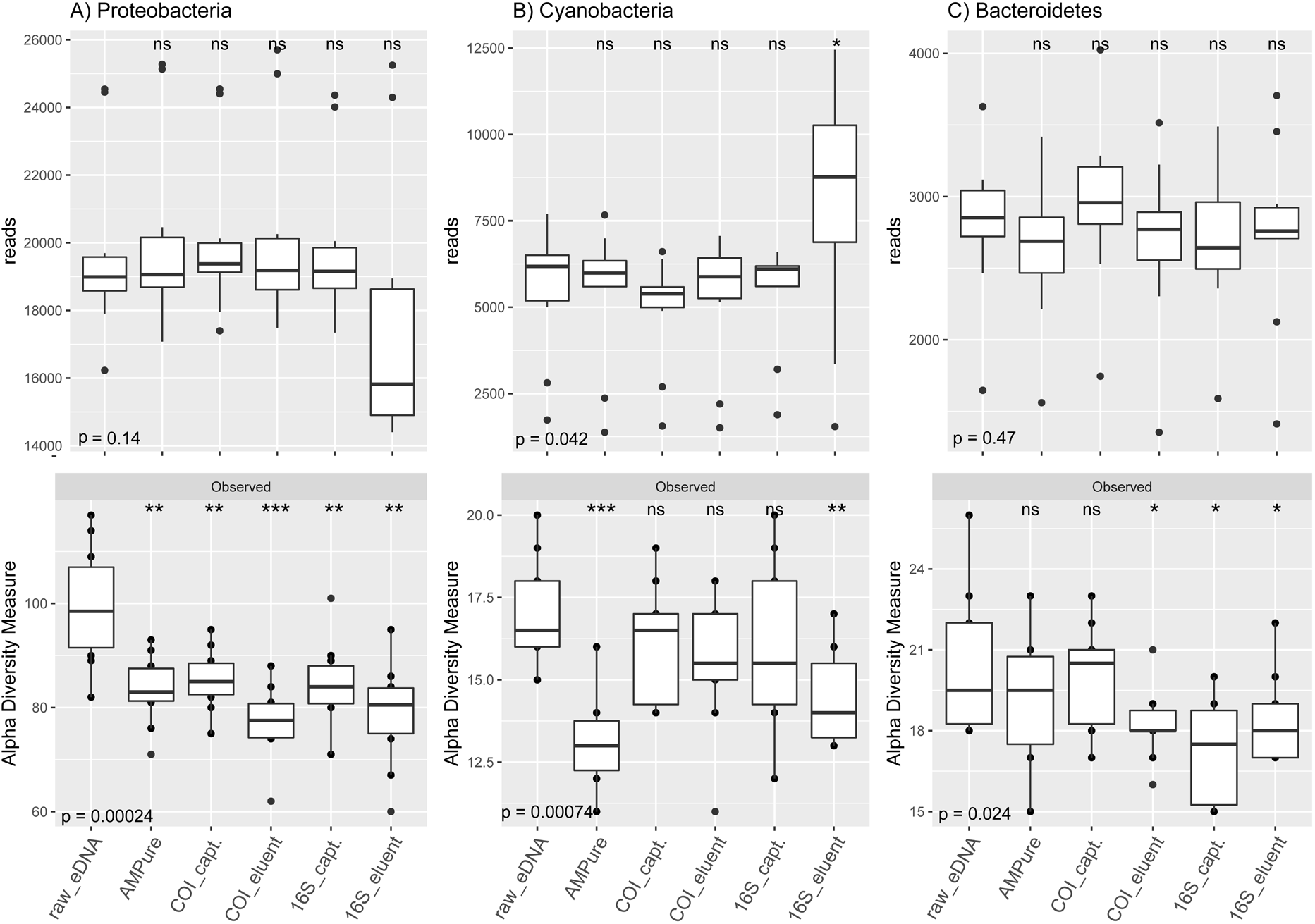
*16S* read abundance and ASV richness. Total reads (top) and ASV richness (bottom) for Protebacteria (A), Cyanobacteria (B) and Bacteroidetes ASVs of the *16S* library. On the x-axis are the different template types used from each step of the targeted capture protocol. The p-values at the bottom show the result of the Kruskal-Wallis test for the whole group, while significant results of Wilcoxon pairwise test between reference group raw_eDNA and all other template types are depicted with asterisks. (Significance levels: * = p < 0.05, ** = p < 0.01, *** = p < 0.001). The boxplot limits stretch over the interquartile range, with the horizontal line signifying the median value, and the whiskers reaching to the upper and lower limits. Outliers are shown as points. NOTE: y-axes do not begin at 0 to accommodate the scale differences for the three phyla and to provide a clearer visualisation.

## TAXONOMIC CONFIDENCE

Overall, the *16S* library showed a high proportion of confident family level assignments across all template types, while the opposite was true for the *COI* library (Fig. 8A). In the *COI* library, the total abundance of family level assignments was significantly increased for both *COI* and *16S* captures (Fig. 8B) but is likely attributable to the increases in bacterial reads overall in the *COI* library for these templates (Fig. 5B). Additionally, the increases in reads confidently assigned to family were not translated into higher richness of good confidence ASVs in the *COI* library (Fig. 8C). In the *16S* library, no changes to the total abundance of good confidence reads were observed, yet the richness of confidently assigned ASVs was significantly reduced in all template types amplified for the *16S* library (Fig. 8B, C).

**Figure 8:**
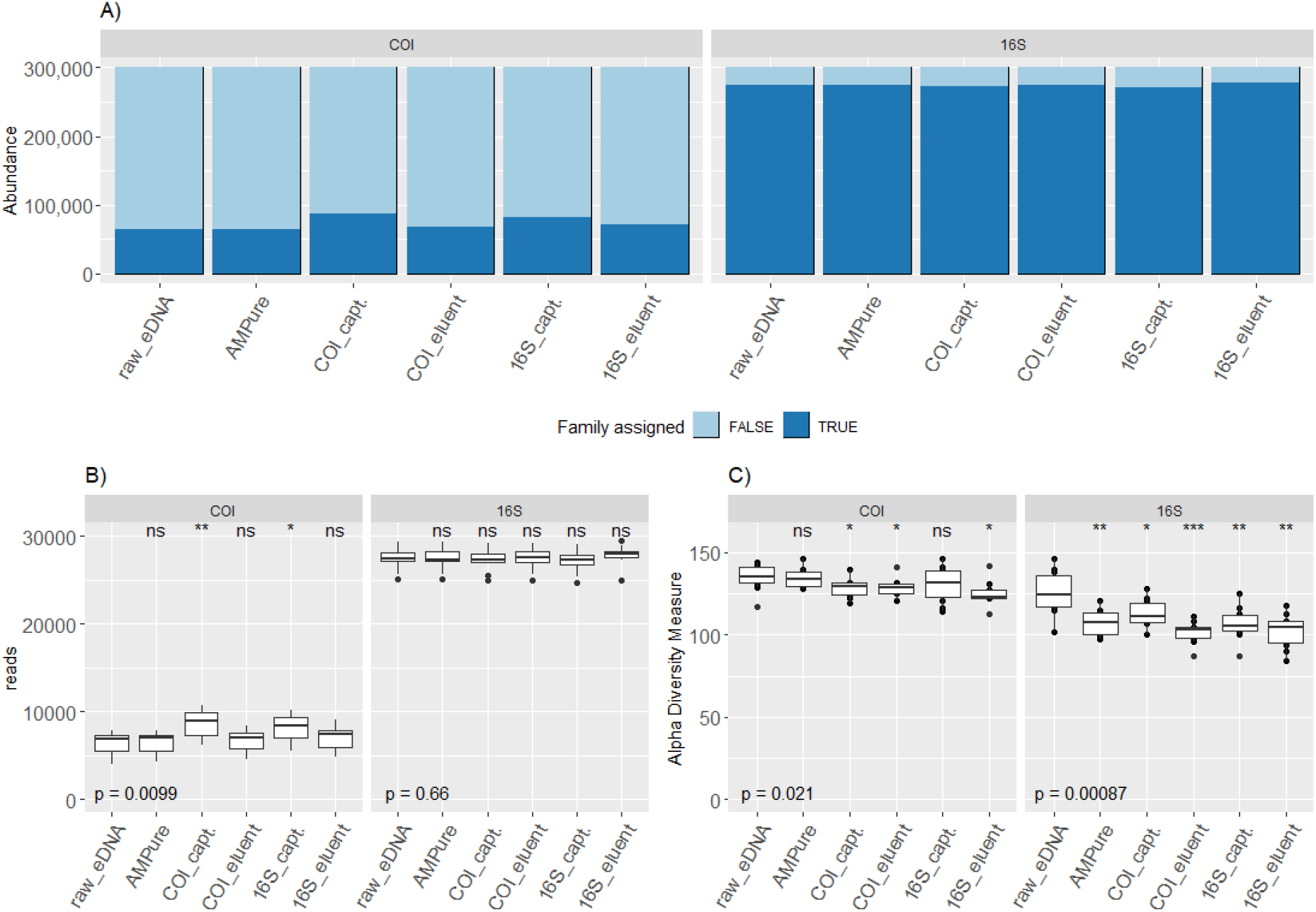
Confident family level assignments of ASVs. A) Proportion of reads that belong to ASVs with high confidence family level assignments. Boxplots show total reads (B) and observed richness (C) of ASVs with high confidence family level assignments. The boxplot limits stretch over the interquartile range, with the horizontal line signifying the median value, and the whiskers reaching to the upper and lower limits. Outliers are shown as points. The p-values at the bottom of boxplots show the resulting p-value of Kruskal-Wallis test for the whole group, while significant results of Wilcoxon pairwise test between reference group raw_eDNA and all other template types are depicted with asterisks (significance levels: * = p < 0.05, ** = p < 0.01, *** = p < 0.001).

## DISCUSSION

This study was done to assess if broadscale metabarcoding primers could act as capture probes for target template enrichment or to remove nontarget templates from eDNA extracts prior to amplification. However, contrary to expectations, the *16S* richness (bacterial) decreased in the remaining DNA pool after using a *COI* probe to isolate target DNA, suggesting that captures with the *COI* probe had a stronger effect on the bacterial template pool than the intended target group of metazoans, and that a significant amount of bacterial DNA was captured with the *COI* probe. Additionally, reductions in metazoan richness were observed in the *COI* capture templates.

In theory, if the captures of bacterial DNA were successful using the *16S* probe, there should be less bacterial DNA available for PCR in the *16S* eluents, and therefore a reduction in relation to the raw eDNA in both bacterial reads and richness could be expected. Here, demonstrating the ability of a capture probe to isolate larger fragments of target DNA, increased availability of *COI*-like bacterial DNA was apparent in the *16S* capture templates based on the increased reads obtained for the kingdom Bacteria when amplifying for *COI* (i.e., the *COI* library). Despite the read counts in the eluents remaining relatively similar to the raw eDNA, richness of bacterial *COI*-like templates was significantly reduced in the *16S* eluents, suggesting most of the bacterial diversity that is amplifiable with *COI* primers was captured with the *16S* probe. The fact that the *16S* library did not show the same pattern of increase with the captures would indicate that the primer pair used for *16S* is able to amplify most of its intended targets even from very diverse samples (i.e., the raw eDNA).

In fact, both probes seemed to capture longer fragments than just their intended target gene from the eDNA samples, as evidenced by the amplification success of bacterial DNA in both the *16S* and *COI* libraries from captured templates. In fact, the *COI* probe appeared to be more efficient in capturing bacterial DNA than metazoan DNA (Fig. 5A, B). This shines a particularly bright spotlight on the inaccuracy of a popular *COI* primer used in metazoan metabarcoding applications, an issue that has not gone unnoticed in recent literature (Deagle et al. 2014, Collins et al. 2019, Zafeiropoulos et al. 2021). Admittedly, the ability of metazoan *COI* primers to amplify microbial DNA has been known for over a decade (Siddall et al. 2009). Although broadscale biodiversity estimates can be obtained when using universal *COI* primers (e.g., Chapters II-III, Grey et al. (2018), Zafeiropoulos et al. (2021)), their bias towards bacterial templates can lead to severe impacts on future metabarcoding efforts, for instance by causing bacterial sequences mislabelled as eukaryotic taxa to be entered into reference databases (Siddall et al. 2009, Mioduchowska et al. 2018). One way to address this nontarget amplification is to develop new primers, however this usually comes with a cost on the universality (Elbrecht and Leese 2017, Marquina et al. 2018, Sultana et al. 2018, Collins et al. 2019, Leese et al. 2021). Therefore, approaches to isolate the target DNA of broad eukaryotic groups from the nontarget DNA (e.g., bacteria and archaea) prior to amplification are more desirable, to maintain the broad applicability of the *COI* marker for metazoans.

It is possible that the reductions observed in richness of metazoan taxonomic groups in the captured templates is a result of the probes pulling out more exclusively DNA of mitochondrial origin, as it would be easier for the probes to attach to and pull out smaller fragments. This is also supported by the increased proportion of target reads that had a lower-level taxonomic assignment after the *COI* capture. Nuclear mitochondrial pseudogenes (numts) have been identified as a potential source of increased species richness estimates (Song et al. 2008), and especially when using denoising rather than OTU clustering methods (Andújar et al. 2021). Here, the higher richness estimates of metazoans in the raw eDNA samples could be a result of numt amplification in addition to the desired mitochondrial *COI* amplification. One way to avoid amplification of nuclear DNA (and therefore numts), is to isolate mitochondrial DNA prior to any further processing by isolating the entire organelles. Isolation of mitochondria can be done for example by differential centrifugation, which has been successfully applied as an enrichment step to avoid the biases arising from PCR on bulk samples of invertebrates, yet this has not yet been tested on eDNA samples (Zhou et al. 2013). That said, current knowledge of the state of eDNA in different environmental conditions is sparse (Mauvisseau et al. 2021), and isolation of full mitochondrial organelles may end up discarding a lot of dissolved or particle bound target DNA from an environmental sample. Bait capture methods on eDNA samples were tested to enrich target DNA, however it is likely the omission of amplification combined with the high dilution factor of target DNA in environmental samples led to reduced species detection rates when the bait capture approach was compared to more conventional PCR based metabarcoding (Wilcox et al. 2018). Hence, the ideal approach would efficiently isolate mitochondrial DNA – or a part of it spanning over the barcoding region – of metazoans to avoid amplification of numts and nontarget taxa, as well as incorporate a universal amplification step to enhance detection rates.

Furthermore, the results also highlighted gaps in some eukaryotic groups in reference sequence databases, as evidenced by the large proportion of ASVs that were annotated as Unassigned by the RDP taxonomic assignment method, being placed within Eukaryota by the Dark mAtteR iNvestigator (DARN). Lack of coverage exists both at the taxonomic level as well as the biogeographic level. For instance, in the Barcode of Life database (BOLD), less than half of Atlantic Iberian coast molluscs, arthropods and polychaetes have a DNA barcode (Leite et al. 2020), while globally the average species representation across all phyla was estimated at approx. 21% (Kvist 2013). However, this relates to coverage in terms of how many species within a phylum have a representative sequence, whereas in some cases, intraspecific coverage may be more important for making accurate taxonomic assignments. Here, most placements of the unassigned ASVs being made to Haptophyta would suggest that the group is not well represented in the reference database used in the RDP assignments, yet not only were they represented, some ASVs were also confidently assigned to species level within Haptophyta by RDP. However, haptophyte taxa can exhibit high levels of intraspecific genetic variation (Medlin et al. 1996), meaning the group would require a higher number of representative references per species to allow for higher levels of confident taxonomic assignments. In cases like this, tools such as DARN that allow for a broader investigation of reads unassignable by conventional assignment methods may help direct future research and barcoding efforts in terms of taxonomic focus.

## CONCLUSIONS

Although the objective of this study to isolate metazoan DNA from environmental samples was not fully fulfilled, the results presented here provide signposts for multiple different avenues of further research. It demonstrated the bacterial bias of a popular *COI* primer with bacterial DNA hitchhiking on the metazoan probe, but in the meantime the results also showed how a relatively simple capture protocol can increase the proportion of taxonomically assignable reads obtained from environmental samples. It may be possible to address the bias issue with better designed primers, but primer design is as heavily reliant on reference databases as is taxonomic assignment of metabarcoding outputs, and restricting metabarcoding output to only those taxa we know now may lead to datasets that cannot be used for temporal analyses in years to come when more taxa have reference sequences. Additionally, because the capture protocol seemed efficient in capturing bacterial DNA along with metazoan templates, a further investigation to the origin of the DNA may be warranted to account for presence or absence of numt amplicons. Based on the results of this study, the use of the tested universal *COI* primer as a capture probe for metazoan taxa cannot be recommended due to the observed loss of richness after the captures. Additionally, little evidence was found that would support the use of the protocol for bacterial enrichment, as the *16S* primers used seemed capable of capturing most of the bacterial diversity from the raw eDNA samples. Nevertheless, the *COI* probe was capable of capturing fragments of bacterial DNA that were amplifiable with the *16S* primer set, providing evidence of relatively large DNA fragments being isolated with the probe. Indeed, perhaps it is possible to find a truly conserved region on metazoan mitochondrial genomes to use as a “capture-region” for mitochondrial DNA, the captured products of which could then be used for conventional *COI* metabarcoding – even with the most degenerate primers.

## CRediT

S.H. → Conceptualization, Formal analysis, Investigation, Methodology, Project administration, Visualization, Writing – original draft, Writing – review and editing

J.E.L.C. → Methodology, Resources, Supervision

J.C. → Conceptualization, Supervision, Writing – review and editing

## ACKNOWLEDGEMENT

We would like to thank Bernard Ball for his support in the laboratory during this experiment, and Haris Zafeiropoulos for his time in discussing the results.

This research was part funded by the Irish Marine Institute as part of the Burrishoole Ecosystem Observatory Network 2020 (BEYOND 2020 PBA/FS/16/02) and the Government of Ireland Postgraduate Scholarship Programme (GOIPG/2018/362). Support for sample collection and logistics was provided by Operation Wallacea Ltd (UK) and Tela Marine Research Centre (Honduras).

## SUPPLEMENTARY FILES

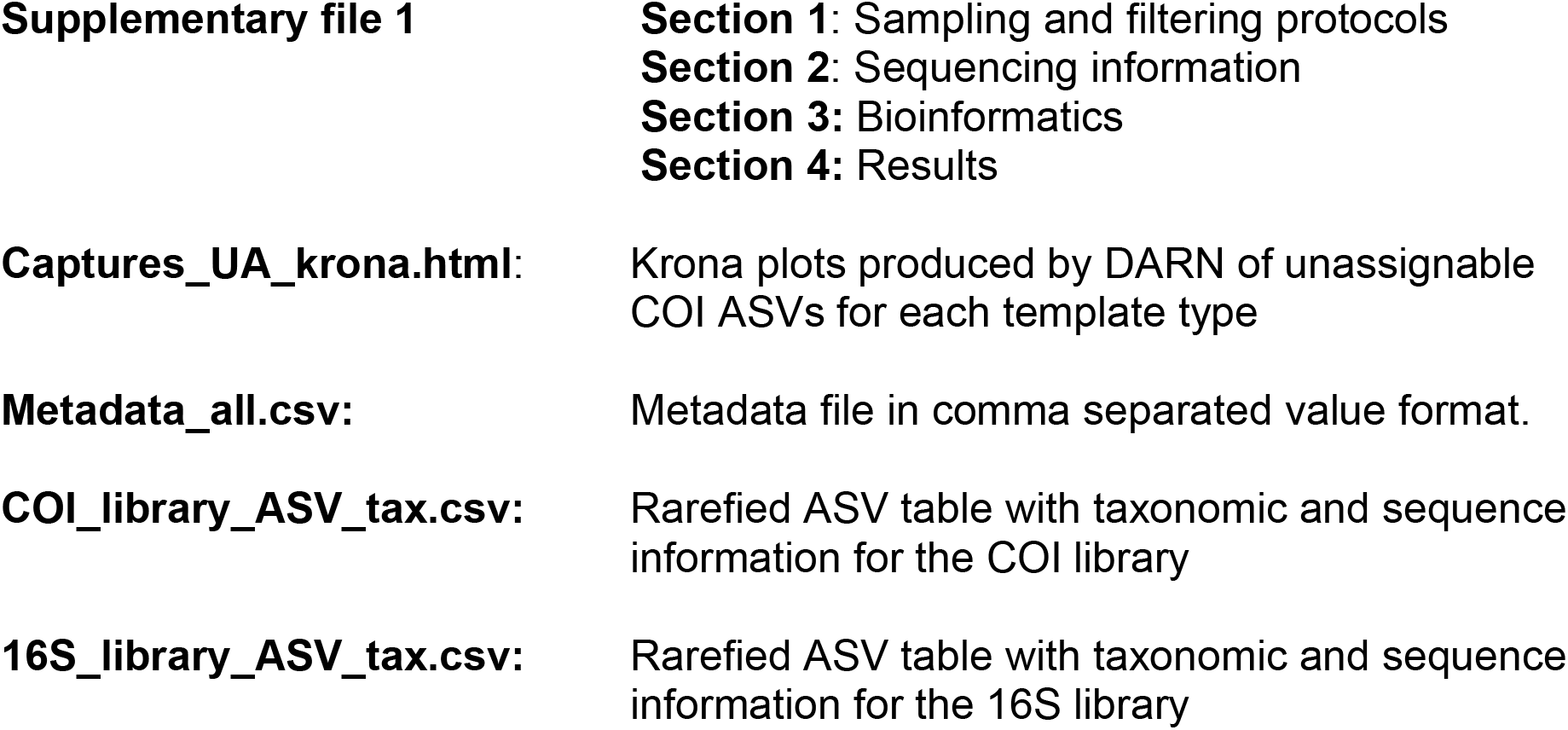

## DATA AVAILABILITY

Raw sequence data available in ENA under project accession PRJEB49001

## Notes

### Competing Interest Statement

The authors have declared no competing interest.

### Summary of Updates

An error was located in the demultiplexing process of the manuscript methods, leading to a re-analysis of the results after correcting the bioinformatics pipeline. The results and discussion have been updated to reflect the newly analysed data. Additionally, the supplementary file ASV_table_with_tax.csv (ASV abundance table with taxonomy) was separated into two files, one for each COI and 16S libraries.

## REFERENCES

Andrews S (2010) FastQC: A Quality Control Tool for High Throughput Sequence Data [Online]. Available from: http://www.bioinformatics.babraham.ac.uk/projects/fastqc/.

Andújar C, Creedy TJ, Arribas P, López H, Salces-Castellano A, Pérez-Delgado AJ, Vogler AP, Emerson BC (2021) Validated removal of nuclear pseudogenes and sequencing artefacts from mitochondrial metabarcode data. Molecular Ecology Resources 21: 1772–1787. https://doi.org/ https://doi.org/10.1111/1755-0998.13337

Antich A, Palacín C, Cebrian E, Golo R, Wangensteen OS, Turon X (2021) Marine biomonitoring with eDNA: Can metabarcoding of water samples cut it as a tool for surveying benthic communities? Molecular Ecology 30: 3175–3188. https://doi.org/ https://doi.org/10.1111/mec.15641

Biggs J, Ewald N, Valentini A, Gaboriaud C, Dejean T, Griffiths RA, Foster J, Wilkinson JW, Arnell A, Brotherton P, Williams P, Dunn F (2015) Using eDNA to develop a national citizen science-based monitoring programme for the great crested newt (Triturus cristatus). Biological Conservation 183: 19–28. https://doi.org/ https://doi.org/10.1016/j.biocon.2014.11.029

Callahan BJ, McMurdie PJ, Rosen MJ, Han AW, Johnson AJA, Holmes SP (2016) DADA2: high-resolution sample inference from Illumina amplicon data. Nature methods 13: 581. https://doi.org/10.1038/nmeth.3869

Cilleros K, Valentini A, Allard L, Dejean T, Etienne R, Grenouillet G, Iribar A, Taberlet P, Vigouroux R, Brosse S (2019) Unlocking biodiversity and conservation studies in high-diversity environments using environmental DNA (eDNA): A test with Guianese freshwater fishes. Molecular Ecology Resources 19: 27–46. https://doi.org/10.1111/1755-0998.12900

Collins RA, Bakker J, Wangensteen OS, Soto AZ, Corrigan L, Sims DW, Genner MJ, Mariani S (2019) Non-specific amplification compromises environmental DNA metabarcoding with COI. Methods in Ecology and Evolution 10: 1985–2001. https://doi.org/10.1111/2041-210X.13276

Deagle BE, Jarman SN, Coissac E, Pompanon F, Taberlet P (2014) DNA metabarcoding and the cytochrome c oxidase subunit I marker: not a perfect match. Biology Letters 10: 20140562. https://doi.org/10.1098/rsbl.2014.0562

Dowle EJ, Pochon X, C. Banks J, Shearer K, Wood SA (2016) Targeted gene enrichment and high-throughput sequencing for environmental biomonitoring: a case study using freshwater macroinvertebrates. Molecular Ecology Resources 16: 1240–1254. https://doi.org/10.1111/1755-0998.12488

Elbrecht V, Leese F (2015) Can DNA-Based Ecosystem Assessments Quantify Species Abundance? Testing Primer Bias and Biomass—Sequence Relationships with an Innovative Metabarcoding Protocol. PLOS ONE 10: e0130324. Available from: https://doi.org/10.1371/journal.pone.0130324.

Elbrecht V, Leese F (2017) Validation and Development of COI Metabarcoding Primers for Freshwater Macroinvertebrate Bioassessment. Frontiers in Environmental Science 5: 11. https://doi.org/10.3389/fenvs.2017.00011

Evans NT, Li Y, Renshaw MA, Olds BP, Deiner K, Turner CR, Jerde CL, Lodge DM, Lamberti GA, Pfrender ME (2017) Fish community assessment with eDNA metabarcoding: effects of sampling design and bioinformatic filtering. Canadian Journal of Fisheries and Aquatic Sciences 74: 1362–1374. https://doi.org/10.1139/cjfas-2016-0306

Ficetola GF, Pansu J, Bonin A, Coissac E, Giguet-Covex C, De Barba M, Gielly L, Lopes CM, Boyer F, Pompanon F, Rayé G, Taberlet P (2014) Replication levels, false presences and the estimation of the presence/absence from eDNA metabarcoding data. Molecular Ecology Resources 15: 543–556. https://doi.org/10.1111/1755-0998.12338

Frøslev TG, Kjøller R, Bruun HH, Ejrnæs R, Brunbjerg AK, Pietroni C, Hansen AJ (2017) Algorithm for post-clustering curation of DNA amplicon data yields reliable biodiversity estimates. Nature Communications 8: 1188. https://doi.org/10.1038/s41467-017-01312-x

Furlan EM, Gleeson D, Hardy CM, Duncan RP (2016) A framework for estimating the sensitivity of eDNA surveys. Molecular Ecology Resources 16: 641–654. https://doi.org/ https://doi.org/10.1111/1755-0998.12483

Gargan LM, Brooks PR, Vye SR, Ironside JE, Jenkins SR, Crowe TP, Carlsson J (2021) The use of environmental DNA metabarcoding and quantitative PCR for molecular detection of marine invasive non-native species associated with artificial structures. Biological Invasions. https://doi.org/10.1007/s10530-021-02672-8

Grey EK, Bernatchez L, Cassey P, Deiner K, Deveney M, Howland KL, Lacoursière-Roussel A, Leong SCY, Li Y, Olds B, Pfrender ME, Prowse TAA, Renshaw MA, Lodge DM (2018) Effects of sampling effort on biodiversity patterns estimated from environmental DNA metabarcoding surveys. Scientific Reports 8: 8843. https://doi.org/10.1038/s41598-018-27048-2

Hajibabaei M, Porter TM, Robinson C V, Baird DJ, Shokralla S, Wright MTG (2019) Watered-down biodiversity? A comparison of metabarcoding results from DNA extracted from matched water and bulk tissue biomonitoring samples. PLOS ONE 14: e0225409. Available from: https://doi.org/10.1371/journal.pone.0225409.

Jeunen G-J, Lamare MD, Knapp M, Spencer HG, Taylor HR, Stat M, Bunce M, Gemmell NJ (2020) Water stratification in the marine biome restricts vertical environmental DNA (eDNA) signal dispersal. Environmental DNA 2: 99–111. https://doi.org/ https://doi.org/10.1002/edn3.49

Kebschull JM, Zador AM (2015) Sources of PCR-induced distortions in high-throughput sequencing data sets. Nucleic Acids Research 43: e143–e143. https://doi.org/10.1093/nar/gkv717

Kvist S (2013) Barcoding in the dark?: A critical view of the sufficiency of zoological DNA barcoding databases and a plea for broader integration of taxonomic knowledge. Molecular Phylogenetics and Evolution 69: 39–45. https://doi.org/ https://doi.org/10.1016/j.ympev.2013.05.012

Leese F, Sander M, Buchner D, Elbrecht V, Haase P, Zizka VMA (2021) Improved freshwater macroinvertebrate detection from environmental DNA through minimized nontarget amplification. Environmental DNA 3: 261–276. https://doi.org/ https://doi.org/10.1002/edn3.177

Leite BR, Vieira PE, Teixeira MAL, Lobo-Arteaga J, Hollatz C, Borges LMS, Duarte S, Troncoso JS, Costa FO (2020) Gap-analysis and annotated reference library for supporting macroinvertebrate metabarcoding in Atlantic Iberia. Regional Studies in Marine Science 36: 101307. https://doi.org/ https://doi.org/10.1016/j.rsma.2020.101307

Leray M, Yang JY, Meyer CP, Mills SC, Agudelo N, Ranwez V, Boehm JT, Machida RJ (2013) A new versatile primer set targeting a short fragment of the mitochondrial COI region for metabarcoding metazoan diversity: application for characterizing coral reef fish gut contents. Frontiers in Zoology 10: 34. https://doi.org/10.1186/1742-9994-10-34

Maggia ME, Vigouroux Y, Renno JF, Duponchelle F, Desmarais E, Nunez J, García-Dávila C, Carvajal-Vallejos FM, Paradis E, Martin JF, Mariac C (2017) DNA Metabarcoding of Amazonian Ichthyoplankton Swarms. PLOS ONE 12: e0170009. Available from: https://doi.org/10.1371/journal.pone.0170009.

Mariac C, Vigouroux Y, Duponchelle F, García-Dávila C, Nunez J, Desmarais E, Renno JF (2018) Metabarcoding by capture using a single COI probe (MCSP) to identify and quantify fish species in ichthyoplankton swarms. PLOS ONE 13: e0202976. Available from: https://doi.org/10.1371/journal.pone.0202976.

Marquina D, Andersson AF, Ronquist F (2018) New mitochondrial primers for metabarcoding of insects, designed and evaluated using in silico methods. Molecular Ecology Resources 0. https://doi.org/10.1111/1755-0998.12942

Martin M (2011) Cutadapt removes adapter sequences from high-throughput sequencing reads. EMBnet. journal 17: pp--10. https://doi.org/10.14806/ej.17.1.200

Mauvisseau Q, Harper L, Sander M, Hanner RH, Kleyer H, Deiner K (2021) The multiple states of environmental {DNA} and what is known about their persistence in aquatic environments. https://doi.org/10.22541/au.163638394.41572509/v1

McMurdie PJ, Holmes S (2013) phyloseq: An R Package for Reproducible Interactive Analysis and Graphics of Microbiome Census Data. PLOS ONE 8: e61217. Available from: https://doi.org/10.1371/journal.pone.0061217.

Medlin LK, Barker GLA, Campbell L, Green JC, Hayes PK, Marie D, Wrieden S, Vaulot D (1996) Genetic characterisation of Emiliania huxleyi (Haptophyta). Journal of Marine Systems 9: 13–31. https://doi.org/ https://doi.org/10.1016/0924-7963(96)00013-9

Mioduchowska M, Czyż MJ, Gołdyn B, Kur J, Sell J (2018) Instances of erroneous DNA barcoding of metazoan invertebrates: Are universal cox1 gene primers too “universal”? PLOS ONE 13: e0199609. Available from: https://doi.org/10.1371/journal.pone.0199609.

O’Donnell JL, Kelly RP, Shelton AO, Samhouri JF, Lowell NC, Williams GD (2017) Spatial distribution of environmental DNA in a nearshore marine habitat. PeerJ 5. https://doi.org/10.7717/peerj.3044

Oksanen J, Kindt R, Legendre P, Hara B, Simpson G, Solymos P, Henry M, Stevens H, Maintainer H, Oksanen@oulu jari (2009) The vegan Package.

Port JA, O’Donnell JL, Romero-Maraccini OC, Leary PR, Litvin SY, Nickols KJ, Yamahara KM, Kelly RP (2016) Assessing vertebrate biodiversity in a kelp forest ecosystem using environmental DNA. Molecular Ecology 25: 527–541. https://doi.org/10.1111/mec.13481

Porter TM, Hajibabaei M (2018) Automated high throughput animal CO1 metabarcode classification. Scientific Reports 8: 4226. https://doi.org/10.1038/s41598-018-22505-4

Sard NM, Herbst SJ, Nathan L, Uhrig G, Kanefsky J, Robinson JD, Scribner KT (2019) Comparison of fish detections, community diversity, and relative abundance using environmental DNA metabarcoding and traditional gears. Environmental DNA 1: 368–384. https://doi.org/ https://doi.org/10.1002/edn3.38

Siddall ME, Fontanella FM, Watson SC, Kvist S, Erséus C (2009) Barcoding Bamboozled by Bacteria: Convergence to Metazoan Mitochondrial Primer Targets by Marine Microbes. Systematic Biology 58: 445–451. https://doi.org/10.1093/sysbio/syp033

Sigsgaard EE, Nielsen IB, Carl H, Krag MA, Knudsen SW, Xing Y, Holm-Hansen TH, Møller PR, Thomsen PF (2017) Seawater environmental DNA reflects seasonality of a coastal fish community. Marine Biology. https://doi.org/10.1007/s00227-017-3147-4

Song H, Buhay JE, Whiting MF, Crandall KA (2008) Many species in one: DNA barcoding overestimates the number of species when nuclear mitochondrial pseudogenes are coamplified. Proceedings of the National Academy of Sciences 105: 13486–13491. https://doi.org/10.1073/pnas.0803076105

Stat M, Huggett MJ, Bernasconi R, DiBattista JD, Berry TE, Newman SJ, Harvey ES, Bunce M (2017) Ecosystem biomonitoring with eDNA: metabarcoding across the tree of life in a tropical marine environment. Scientific Reports 7: 12240. https://doi.org/10.1038/s41598-017-12501-5

Sultana S, Ali ME, Hossain MAAM, Asing, Naquiah N, Zaidul ISM (2018) Universal mini COI barcode for the identification of fish species in processed products. Food Research International 105: 19–28. https://doi.org/10.1016/j.foodres.2017.10.065

Thomsen PF, Kielgast J, Iversen LL, Møller PR, Rasmussen M, Willerslev E (2012) Detection of a Diverse Marine Fish Fauna Using Environmental DNA from Seawater Samples. PLOS ONE 7: e41732. Available from: https://doi.org/10.1371/journal.pone.0041732.

Valentini A, Taberlet P, Miaud C, Civade R, Herder J, Thomsen PF, Bellemain E, Besnard A, Coissac E, Boyer F, Gaboriaud C, Jean P, Poulet N, Roset N, Copp GH, Geniez P, Pont D, Argillier C, Baudoin J-M, Peroux T, Crivelli AJ, Olivier A, Acqueberge M, Le Brun M, Møller PR, Willerslev E, Dejean T (2016) Next-generation monitoring of aquatic biodiversity using environmental DNA metabarcoding. Molecular Ecology 25: 929–942. https://doi.org/ https://doi.org/10.1111/mec.13428

Wang Q, Garrity GM, Tiedje JM, Cole JR (2007) Naïve Bayesian Classifier for Rapid Assignment of rRNA Sequences into the New Bacterial Taxonomy. Applied and Environmental Microbiology 73: 5261 LP – 5267. https://doi.org/10.1128/AEM.00062-07

Wilcox TM, Zarn KE, Piggott MP, Young MK, McKelvey KS, Schwartz MK (2018) Capture enrichment of aquatic environmental DNA: A first proof of concept. Molecular Ecology Resources 18: 1392–1401. https://doi.org/10.1111/1755-0998.12928

Zafeiropoulos H, Gargan L, Hintikka S, Pavloudi C, Carlsson J (2021) The Dark mAtteR iNvestigator (DARN) tool: getting to know the known unknowns in COI amplicon data. Metabarcoding and Metagenomics 5: e69657. Available from: https://doi.org/10.3897/mbmg.5.69657.

Zhou X, Li Y, Liu S, Yang Q, Su X, Zhou L, Tang M, Fu R, Li J, Huang Q (2013) Ultra-deep sequencing enables high-fidelity recovery of biodiversity for bulk arthropod samples without PCR amplification. GigaScience 2: 4. https://doi.org/10.1186/2047-217X-2-4

