## Supplementary material for "The bacterial hitchhiker’s guide to COI: Universal primer-based COI capture probes fail to exclude bacterial DNA, but 16S capture leaves metazoa behind": Captures_UA_krona.html

Javascript must be enabled to view this page.

magnitude
magnitudeUnassigned

darn\_counts\_raw\_eDNA\_krona
darn\_counts\_AMPure\_krona
darn\_counts\_COI\_capt\_krona
darn\_counts\_COI\_eluent\_krona
darn\_counts\_16S\_capt\_krona
darn\_counts\_16S\_eluent\_krona

840833794824803782

102131
179179187183192177

000000
177176183181187174

177176183181187174
545152514847

262627292724
000000

000000
262627292724

000000
262627292724

262627292724
000000

252526282623
262627292724

000000
111111

111111
000000

000000
111111

111111

000000
9799104101112103

9799104101112103
000000

9799104101112103

132122
000000

000000
132122

132122
000000

000000
122122

122122
000000

122122
000000

122122
000000

000000
122122

000000
122122

122122
000000

000000
122122

000000
122122

000000
122122

122122
000000

122122

0
1

1

411311

169173170178171169
657653606638610604

000000
632345

632345
000000

632345

887888
111111

443444
000000

443444

222222

111111

111111
000000

111111
000000

111111

10111110911
000000

10111110911
324223

675666

122212

121211
000000

121111
121211

0
1

1
0

1
0

1

000
111

11
00

11

1
0

1
0

0
1

0
1

0
1

1

000000
212124242124

181922231822
212124242124

322132
000000

322132
000000

322132

118109109
101111

2213

887768

000000
161413151511

161413151511
151312141411

11111
00000

11111
00000

00000
11111

11111

553534
495042514547

111111

00000
22121

00000
22121

22121

000000
151614161514

111211111111
000000

111211111111
000000

000000
111211111111

111211111111

0000
1121

0000
1121

0000
1121

1121

333333

00000
11112

11112
00000

11112
00000

11112
00000

11112
00000

11112

000000
252523262525

11111

242422262424

112333
444666

000000
332333

332333

000000
131414131414

131414131414
000000

131414131414

000000
333333

222222
333333

111111
000000

111111
000000

111111
000000

111111

333333
000000

333333
000000

333333

876967
000000

554645
876967

322322

000000
544445

544445

2222
0000

2222
0000

0000
2222

0000
2222

2222
0000

2222

000000
111111

111111

111111
000000

000000
111111

000000
111111

000000
111111

111111
000000

111111
000000

111111

300298263273264253
000000

143137124132121115
300298263273264253

10510489899589
000000

000000
10510489899589

10510489899589

1

515750524849
000000

000000
515750524849

515750524849

0
1

0
1

1

000000
112122

112122

321211

191919171618
000000

000000
191919171618

191919171618
000000

191919171618
000000

000000
191919171618

191919171618

111111
000000

111111

000000
111211

111211
000000

111111

0
1

1
