## Supplementary material for "The bacterial hitchhiker’s guide to COI: Universal primer-based COI capture probes fail to exclude bacterial DNA, but 16S capture leaves metazoa behind": Supplementary_File_1.docx

The bacterial hitchhiker’s guide to COI: Universal primer-based COI capture probes fail to exclude bacterial DNA, but 16S capture leaves metazoa behind

Authors: Sanni Hintikka, Jeanette Carlsson, Jens Carlsson

**Section 1 - Sampling and Filtering protocols**

### Site description and sampling protocol

#### Tela – Honduras

The study area of Tela Bay (15°46'42.4"N 87°28'23.7"W), on the north coast of Honduras, has a tropical rainforest climate (no dry season, increased rainfall between October and December) with an offshore coral reef (Banco Capiro, 15°51'48.6"N 87°29'42.9"W). The coral reef is part of the Meso-American Barrier Reef (second largest barrier reef system in the world) and lies approximately 8km north from the coast of Tela, a town with a population of ca. 38K people. We used 5 eDNA samples from two subsites on the reef each, for a total of 10 eDNA samples. The samples were collected using a submersible Ruttner sampler, which was sterilised with 10% bleach between each sampling point. The water collected by the Ruttner sampler was dispensed into re-usable 1L sampling bottles. To minimise contamination, these re-usable sampling bottles were sterilised prior to sampling by submersing them in 10% bleach for a minimum of 20 minutes, followed by a tap water rinse and placing them in the sun (caps closed) for natural UV exposure for a minimum of 20 minutes.

### Filtering protocol

Filtering of water samples was performed in a dedicated room onsite. Only persons directly involved in the filtering process were allowed into the filtering room during the process to minimise risk of contamination. Filtering equipment (filter funnels, trays, tweezers, etc.) were sterilised using a similar approach, with a 10% bleach bath for 20 min, followed by rinse with tap water and dry in an access-controlled room. The filter funnels were assembled using sterilised equipment (trays and tweezers), stored in new clean ziplock bags and placed in the sun for UV exposure for a minimum 20 min. For each ecological replicate collected, 1L of water was filtered through a Whatman® glass microfiber filter (Grade 934-AH®, pore size 1.5 µm), and stored in a 2mL screwcap tube with silica beads to help remove any remaining moisture that could accelerate DNA degradation. If a filter got saturated before 1L had been filtered through, a second filter was used for the remaining water, and the amount of water filtered through each was noted. The tubes containing the eDNA filters were stored out of sunlight at -20 ˚C until transportation to University College Dublin, Ireland, for laboratory processing.

**Section 2 - Sequence information**

### Sample identifier tags

Table S1: Tags used in 5’ end of forward and reverse primers. The tags were the same for forward and reverse primers, with the same numbering order (i.e., f1 = 5’-TAG1-FWD_primer-3’; r1 = 5’-TAG1-REV_primer-3’)

| TAG | TAG_NR |
| --- | --- |
| AGACGC | 1 |
| AGTGTA | 2 |
| ACTAGC | 3 |
| ACAGTC | 4 |
| ATCGAC | 5 |
| ATGTCG | 6 |
| CTCTAG | 7 |
| CATCAC | 8 |
| TACGAG | 9 |
| ACTCTG | 10 |

**Section 3 - Bioinformatics**

Demultiplexing with *cutadapt*

COI library:

$ cutadapt \

-e 0.08 --no-indels -g file:fwd.fasta -G file:rev.fasta \

-o COI/1_demux/{name1}-{name2}_1.fastq \

-p COI/1_demux/{name1}-{name2}_2.fastq \

COI_R1.fastq COI_R2.fastq --minimum-length 100 > COI/fwd_orient.cutadapt.stat

$ cutadapt \

-e 0.08 --no-indels -G file:fwd.fasta -g file:rev.fasta \

-o COI/2_demux/{name1}-{name2}_1.fastq \

-p COI/2_demux/{name1}-{name2}_2.fastq \

COI_R1.fastq COI_R2.fastq --minimum-length 100 > COI/rev_orient.cutadapt.stat

16S library:

$ cutadapt \

-e 0.08 --no-indels -g file:fwd_bac.fasta -G file:rev_bac.fasta \

-o BAC/1_demux/{name1}-{name2}_1.fastq \

-p BAC/1_demux/{name1}-{name2}_2.fastq \

BAC_R1.fastq BAC_R2.fastq --minimum-length 100 > BAC/fwd_orient.cutadapt.stat

$ cutadapt \

-e 0.08 --no-indels -G file:fwd_bac.fasta -g file:rev_bac.fasta \

-o BAC/2_demux/{name1}-{name2}_1.fastq \

-p BAC/2_demux/{name1}-{name2}_2.fastq \

BAC_R1.fastq BAC_R2.fastq --minimum-length 100 > BAC/rev_orient.cutadapt.stat

The resulting .fastq files that matched used tag combinations were renamed with the corresponding sample names using *multiple move* command *mmv* with a .txt sample index file:

sample index file format:

f1-r1_[12].fastq	matched/A1_#1.fastq

f1-r2_[12].fastq	matched/A2_#1.fastq

f1-r3_[12].fastq	matched/A3_#1.fastq

…

**Section 4 - Results**

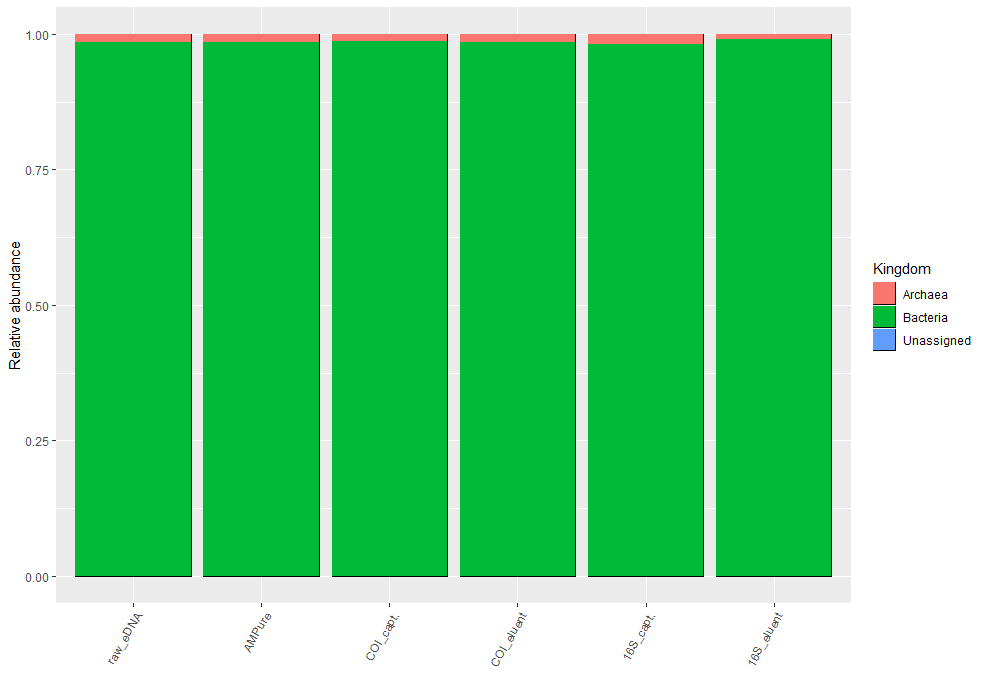

Figure S1: Relative abundance of kingdoms detected in the 16S library.

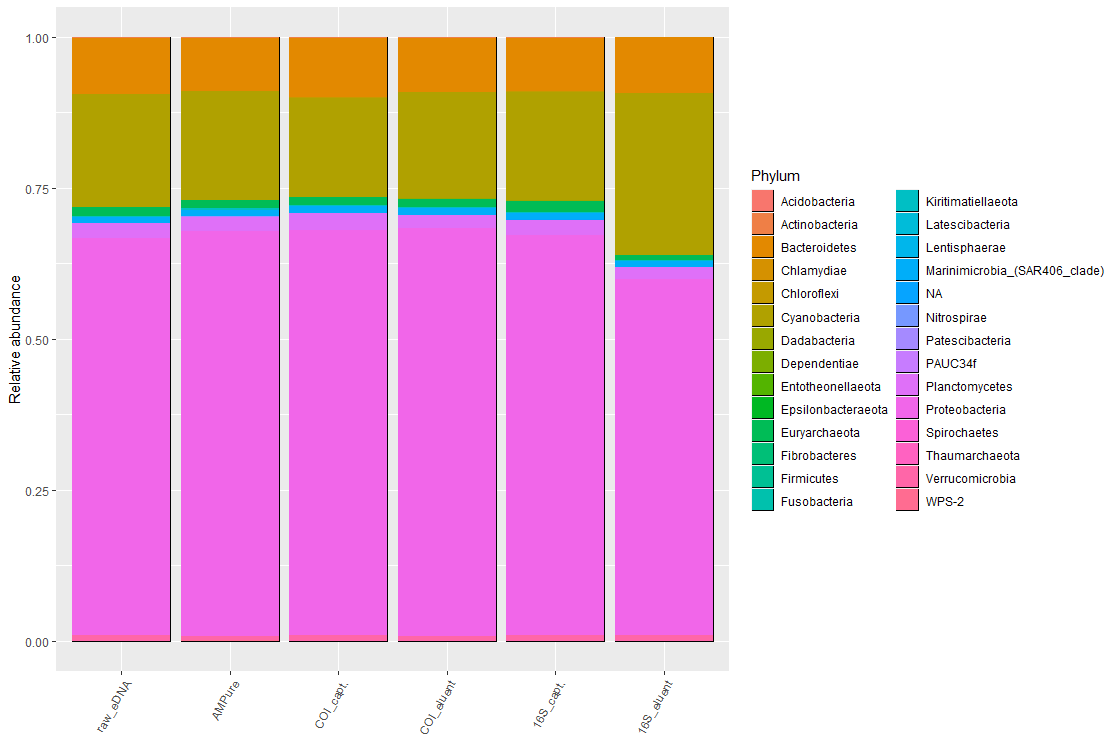

Figure S2: Relative abundance of phyla detected in the 16S library.
